# Comparative analysis of the plasma metabolome of migrating passerines during stopover: Novel insights into flight metabolism

**DOI:** 10.1101/2024.01.09.574878

**Authors:** By Adi Domer, Weronika Jasinska, Leah Rosental, Eyal Shochat, Saleh Alseekh, Alisdair R. Fernie, Yariv Brotman, Ofer Ovadia

## Abstract

During long distance migration, many birds may experience periods of either prolonged fasting, during endurance flights, or extensive feeding during stopovers. It was previously shown that habitat selection during stopover can largely affect the migration outcome of an individual. Despite decades of research of the avian metabolism during stopover and migration, many questions have remained unanswered, as such research mainly focused on targeted metabolites and fat metabolism. Here, we examined the plasma-metabolome of migrating passerines prior to their crossing the Sahara Desert. Birds were sampled at two sites populated by Pistacia trees, bearing fat-rich fruits, and at an additional site dominated by blooming Eucalyptus trees. The blood samples were analyzed using both GC-MS and LC-MS, using an untargeted approach. We found that birds from one of the sites had a distinguish metabolic profile, suggesting recent landing. Examination of metabolic pathways activated during stopovers indicated a crucial role for cycling glucose through the Cori and Cahill cycles in resting and recovery processes. This novel perspective, conducted on free-ranging birds, suggests the evolution of avian insulin resistance due to factors such as endurance exercise, fasting, and a preference for fatty acid oxidation during migration, akin to cell trauma recovery. Additionally, we investigated inter-site variations in birds’ metabolic profiles. Significant variations were observed in both polar and lipophilic metabolites among the sites. Differences in polar metabolites were primarily attributed to variations in the physiological state of the birds between sites, while distinctions in the lipophilic profiles of rested birds were linked to variations in their primary food sources. This study underscores the challenge of interpreting commonly used indicators for assessing migrating birds’ physiological states and site quality, which are predominantly derived from lipid metabolism, in complex ecological systems.

## Introduction

Animal migration – one of nature’s most visible and widespread phenomena (Wilcove and Wikelski 2008) – has evolved independently several in varying taxa (Aidley 1981). Migratory behavior is widely common in the avian taxon, with approximately half of the species performing some type of migratory movements (Berthold 1996). Migratory birds alternate between two extreme physiological states, fasting during the long-distance endurance flights and resting or extensively feeding during stopovers (McWilliams and Karasov 2005). Hence, selecting a proper stopover site is crucial for long-distance migrants as low fuel deposition rates can extend their total migration period and affect their fitness (Gómez et al. 2017, Domer et al. 2021). We present an in-depth comparative analysis of the untargeted metabolomic profiling of wild migratory passerines sampled in the eastern Mediterranean region during autumn, along one of the most important flyways in the old world. Previous targeted metabolic studies on wild and captive migratory birds have provided important insights into flight metabolism modalities, including fuel utilization (Jenni and Jenni-Eiermann 1998, Jenni-Eiermann et al. 2002, Smith et al. 2007), protein catabolism (Robin et al. 1987, Smith et al. 2007), and oxidative damage repair of flight muscles (Costantini et al. 2007). While these studies have laid the foundations for the metabolic migration framework, they considered only a few targeted metabolites (Jenni-Eiermann and Jenni 1991, Jenni and Jenni-Eiermann 1998, Jenni-Eiermann et al. 2002, Guglielmo et al. 2005, Seaman et al. 2005) while mainly focusing on lipid metabolism. Flight metabolism comprises many inter-dependent pathways and modalities, some of which have recently gained attention (Levin et al. 2017, Potter et al. 2021, Satoh 2021). To broaden the current perspective of flight metabolism and to better link the different metabolic pathways it comprises, we have adopted an untargeted metabolomic approach.

Within the adopted untargeted approach, we focused on the following metabolic pathways: (1) lipid metabolism, (2) amino acid metabolism, and (3) glucose metabolism.

1. Lipids are considered the primary energy source during endurance migratory flights (Blem 1976, Stevens 2004), accumulated before the migration journey. Plasma triglycerides (TAGs) are usually elevated during refueling (Jenni-Eiermann and Jenni 1992) but may also increase during flight (Bordel and Haase 1993, Schwilch et al. 1996a). Such TAGs differ in their fatty acid (FA) composition in terms of the length of the carbon chain and the unsaturation levels. Most lipid reserves in migrating birds are polyunsaturated FA (PUFA) and they are usually considered as the preferred fuel for endurance exercise (Maillet and Weber 2006). Two additional metabolites reflecting the physiological state of an individual bird are plasma glycerol, which increases during fasting due to high rates of lipolysis (Jenni-Eiermann and Jenni 1991), and plasma β-Hydroxybutyric acid (BUTY), which increases during fasting owing to ketone formation. The level of BUTY increases shortly after exercise (∼20 minutes), indicating post-exercise ketosis that lasts for several hours (Jenni-Eiermann and Jenni 2001). BUTY levels gradually decrease after sufficient rest (∼10 hours). Metabolic studies also highlight birds’ tolerance to hypoxia, which is indicated by elevated plasma lactate (Faraci 1991), as well as by the post-flight metabolic state, during which birds continue lipolysis at a reduced level to meet the energy demands of resting (Jenni-Eiermann 2017).
2. The role of protein catabolism in bird migration was thoroughly investigated (Bauchinger and McWilliams 2012). During long-distance flights, birds catabolize not only lipids but also proteins. These proteins originate in the muscles and other internal organs, especially digestive organs (McWilliams and Karasov 2001, Bauchinger and McWilliams 2012). Free amino acids derived from protein catabolism were previously suggested to serve as substrates for a) gluconeogenesis necessary to meet the brain energy requirements, b) building new energy stores when the fat stores are depleted, and c) maintain water balance during nonstop flights (Gerson and Guglielmo 2011). Additionally, catabolizing protein is known to have antioxidative capacity benefits, as amino acids’ bioactive properties are liberated during detaching from the parent protein in which these peptides are usually inactive (Dai et al. 2017).
3. An additional metabolite of high importance in birds is glucose. Birds are naturally hyperglycemic, maintaining approximately twice the plasma-glucose concentration of mammals at equivalent size while using mechanisms of insulin resistance (Braun and Sweazea 2008). Although the ultimate causation of this phenomenon is largely unknown, recent studies have suggested that hyperglycemia and insulin resistance are related to the drop of oxygen concentrations in the atmosphere at the Permian–Triassic (PT) boundary, forcing theropods to lose certain genes to maximize their efficiency of oxygen usage (Satoh 2021). Indeed, omentin and insulin-sensitive glucose transporter 4 (GLUT4) are considered missing or unfunctional in the bird genome (Braun and Sweazea 2008, Luo et al. 2023). Because these gene products play essential roles in maintaining insulin sensitivity, this loss probably forced theropods to become insulin resistant (Satoh 2021). These high blood glucose levels were also suggested to be correlated with the high metabolic rate and body temperature of birds associated with the extreme energetic requirements of powered flights (Clarke and Rothery 2008, Clarke and Pörtner 2010). Blood glucose levels usually increase after endurance flight (Viswanathan et al. 1987, Schwilch et al. 1996b, Abdel-Rachied et al. 2014), Yet it is not clear if this hyperglycemia represents an adaptive metabolic mechanism or constraint.

We quantified the plasma metabolome of two most common migratory warbler species in Israel: The Eurasian Blackcap (*Sylvia atricapilla*) and Lesser Whitethroat (*Curruca curruca*). Although related, these two species differ in their breeding areas and habitat selection during a stopover. Differences are also manifested in their diet preferences during migration, as the Eurasian Blackcap is more restricted to water consumption (Sapir et al. 2004). Birds were sampled at two previously studied stopover sites dominated by Pistacia trees, bearing fat-rich fruits during autumn (Domer et al. 2018), namely Midreshet Ben-Gurion (hereafter BGS) and Ein-Rimon (hereafter ER), located in arid and semi-arid areas, respectively. While ER is a planted homogeneous *Pistacia atlantica* grove, BGS is a mixed pistachio grove comprising four primary species: *Pistacia atlantica*, *Pistacia chinensis*, *Pistacia vera*, and *Pistacia lentiscus*. Birds were also sampled at a third site, located in the semi-arid area of Israel, ∼11 km south of ER, and mainly populated by autumn-blooming Eucalyptus trees (Negev Brigade Monument, hereafter AN, (31⁰16’N 34⁰49’E)). Previous research showed that fuel accumulation and recapture rates were substantially lower in BGS compared with ER (Domer et al. 2018). These findings may suggest that most birds captured at BGS (arid region) are leaving soon after arrival, and are captured several hours after landing, and most of those caught at ER (semi-arid area) are at a resting/refueling state. Therefore, we hypothesized that plasma metabolome varies among sites, depending on the type of the primary food source (fat-rich fruits vs. nectar) and the physiological state of staging birds (either well rested or landed during the previous night).

## Methods

We conducted metabolomic profiling of the Eurasian Blackcap (N=43, Table 1) and the Lesser Whitethroat (N=30, Table 1). Birds were captured for 3 hours during the morning using mist-nets, opened at first light for three hours. Captured birds were individually tagged with numbered aluminum leg rings, weighed to ±0.1 g with a digital balance, and measured for wing length. Soon after (a few minutes after capture), a blood sample of 0.1ml was extracted from the bird’s jugular vein using 25G insulin needle and heparinized tubes. The blood samples were then stored on ice for several hours, before being centrifuged at 10,000 RPM for 10 min at 4°c. Extracted plasma was maintained at -80°c until processing for metabolomic analyses.

**Table 1.** Number of Blackcaps and Lesser Whitethroats sampled at each study site.

|  | Lesser Whitethroat | Blackcap |
| --- | --- | --- |
| <u>Site</u> |  |  |
| AN | 8 | 5 |
| ER | 11 | 19 |
| BGS | 11 | 19 |

### Metabolomic analyses

#### Lipid and Polar Metabolite Extraction Protocol

Metabolites were extracted from 50 μl of plasma using a protocol described by Hummel et al. (2011). In brief, metabolites from each aliquot were extracted with 1 ml of pre-cooled (−20°C) extraction buffer (homogenous methanol/methyl-*tert*-butyl-ether [1:3] mixture). After 10 min incubation at 4°C and sonication for 10 min in a sonic bath, 500 μl of methanol/water [1:3] mixture was added. Samples were then centrifuged (5 min, 14 000 *g*), leading to a lipophilic and polar phase forming. Five hundred microliters of the lipophilic (upper) phase and 150 μl of the polar phase were collected and dried under a vacuum. The lipophilic phase was resuspended in 200 μl of ACN/isopropanol and used for lipid analysis. The polar phase residue was derivatized for 120 min at 37°C (in 50 μl of 20 mg ml−1 methoxyamine hydrochloride in pyridine) followed by a 30-min treatment at 37°C with 50 μl of MSTFA (with fatty acid methyl esters) and was used for gas chromatography–mass spectrometry (GC–MS) analysis.

#### Lipid Profiling

Samples were processed using UPLC-FT-MS (Hummel et al. 2011) on a C8 reverse-phase column (100 × 2.1 mm × 1.7 μm particle size, Waters) at 60°C. The mobile phases consisted of 1% 1 M NH_4_OAc and 0.1% acetic acid in water (buffer A), and acetonitrile/isopropanol (7:3, UPLC grade BioSolve) supplemented with 1 M NH4Ac and 0.1% acetic acid (buffer B). The following gradient profile was applied: 1 min 45% A, 3 min linear gradient from 45% A to 35% A, 8 min linear gradient from 25% to 11% A, 3 min linear gradient from 11% to 1% A. Finally, after washing the column for 3 min with 1% A the buffer was set back to 45% A and the column was re-equilibrated for 4 min, leading to a total run time of 22 min. The flow rate of the mobile phase was 400 μl/min.

The mass spectra were acquired using a Q-Exactive mass spectrometer (Thermo Fisher, http://www.thermofisher.com) equipped with an ESI interface. All the spectra were recorded using altering full-scan mode, covering a mass range from 150–1500 *m*/*z* at a capillary voltage of 3.0 kV, with a sheath gas flow value of 60 and an auxiliary gas flow of 35. The resolution was set to 30000 with 3 scans per second, restricting the Orbitrap loading time to a maximum of 100 ms with a target value of 1E6 ions. The capillary temperature was set to 150°C, while the drying gas in the heated electrospray source was set to 350°C. The skimmer voltage was held at 25 V while the tube lens was set to a value of 130 V. The spectra were recorded from minute 1 to minute 20 of the UPLC gradients.

Processing of chromatograms, peak detection, and integration was performed using REFINER MS 14.0 (GeneData, http://www.genedata.com) or Xcalibur (Version 3.1, Thermo Fisher, Bremen, Germany). In the first approach, the molecular masses, retention time, and associated peak intensities of the sample were extracted from the raw files, which contained the full-scan MS. Processing MS data included removing the fragmentation information, isotopic peaks, and chemical noise. Further peak filtering on the manually extracted spectra or the aligned data matrices was performed. Obtained features (*m*/*z* at a certain retention time) were queried against an in-house lipid database (Lapidot-Cohen et al. 2020).

#### Polar Metabolite Analysis

The GC–MS system was a gas chromatograph coupled to a time-of-flight mass spectrometer (Pegasus III, Leco). An autosampler system (PAL) injected the samples. Helium was used as carrier gas at a constant flow rate of 2 ml s−1, and gas chromatography was done on a 30-m DB-35 column. The injection temperature was 230°C, and the transfer line and ion source were set to 250°C. The initial temperature of the oven (85°C) increased at a rate of 15°C min−1 up to a final temperature of 360°C. After a solvent delay of 180 s, mass spectra were recorded at 20 scans s−1 with *m*/*z* 70–600 scanning range. Chromatograms and mass spectra were evaluated by using Chroma TOF 1.0 (Leco) (Schauer et al. 2008) together with TargetSearch (Cuadros-Inostroza et al. 2009) and Xcalibur Software (Thermo Scientific). Data for the lipid and polar metabolites is available at Dryad (Domer Adi 2023).

### Statistical analyses

To test for differences in the plasma metabolite composition between birds sampled at the different stopover sites, we used non-metric multidimensional scaling (nMDS) ordinations of the Bray-Curtis dissimilarity matrix, followed by PERMANOVA and SIMPER analyses. The latter allowed quantifying the contribution of different metabolites to the observed inter-site variation. To test for differences in plasma BUTY and TAG levels, we used a generalized linear model (glm) with normal distribution for each response variable, using the site as a categorical variable, and body condition (derived from the residuals of regressing individuals’ body mass against wing length) as a covariate. To test for differences in the levels of specific metabolites (annotated amino acids and non-annotated metabolites detected by SIMPER analyses) among sites, we used multivariate analysis of variance (MANOVA), with annotated metabolites as response variables, the site as a categorical variable, and body condition (derived from the residuals of regressing individuals’ body mass against wing length) as a covariate. All statistical analyses were performed in R 3.4.4 (Team 2013).

## Results

### Polar metabolites

Bird plasma samples were analyzed using GC-MS, generating 414 distinct metabolites. To test for inter-site differences in the composition of these metabolites, we used a non-metric multidimensional scaling (nMDS) ordination of the Bray-Curtis dissimilarity matrix, followed by a PERMANOVA and SIMPER analysis. In both warbler species, the composition of blood polar metabolites varied significantly among sites (Fig. 1; PERMANOVAs: F_2,37_=36.135, P<0.001, R^2^=0.629 and F_2,27_=15.653, P<0.001, R^2^=0.504 for Eurasian Blackcap and Lesser Whitethroat, respectively). The polar metabolic profile of Eurasian Blackcap varied significantly with body condition, derived from the residuals of regressing body mass against wing length (F_1,37_=3.872, P=0.038, R^2^=0.034) but not that of Lesser Whitethroat (F_1,27_=0.563, P=0.567, R^2^=0.009). In both species, the interaction between site and body condition was not significant (F_2,37_= 0.814, P=0.496, R^2^=0.014 and F_2,27_=1.617, P=0.179, R^2^=0.052 for Eurasian Blackcap and Lesser Whitethroat, respectively). Pairwise comparisons revealed that the polar metabolic profile, characterizing birds in BGS varied significantly from that of birds in ER (P=0.003 and P=0.003 for Eurasian Blackcap and Lesser Whitethroat, respectively) and AN (P=0.003 and P=0.003 for Eurasian Blackcap and Lesser Whitethroat, respectively), but not between ER and AN (P=0.078 and P=0.063 for Eurasian Blackcap and Lesser Whitethroat, respectively), although both might be considered as marginally non-significant.

**Figure 1:**
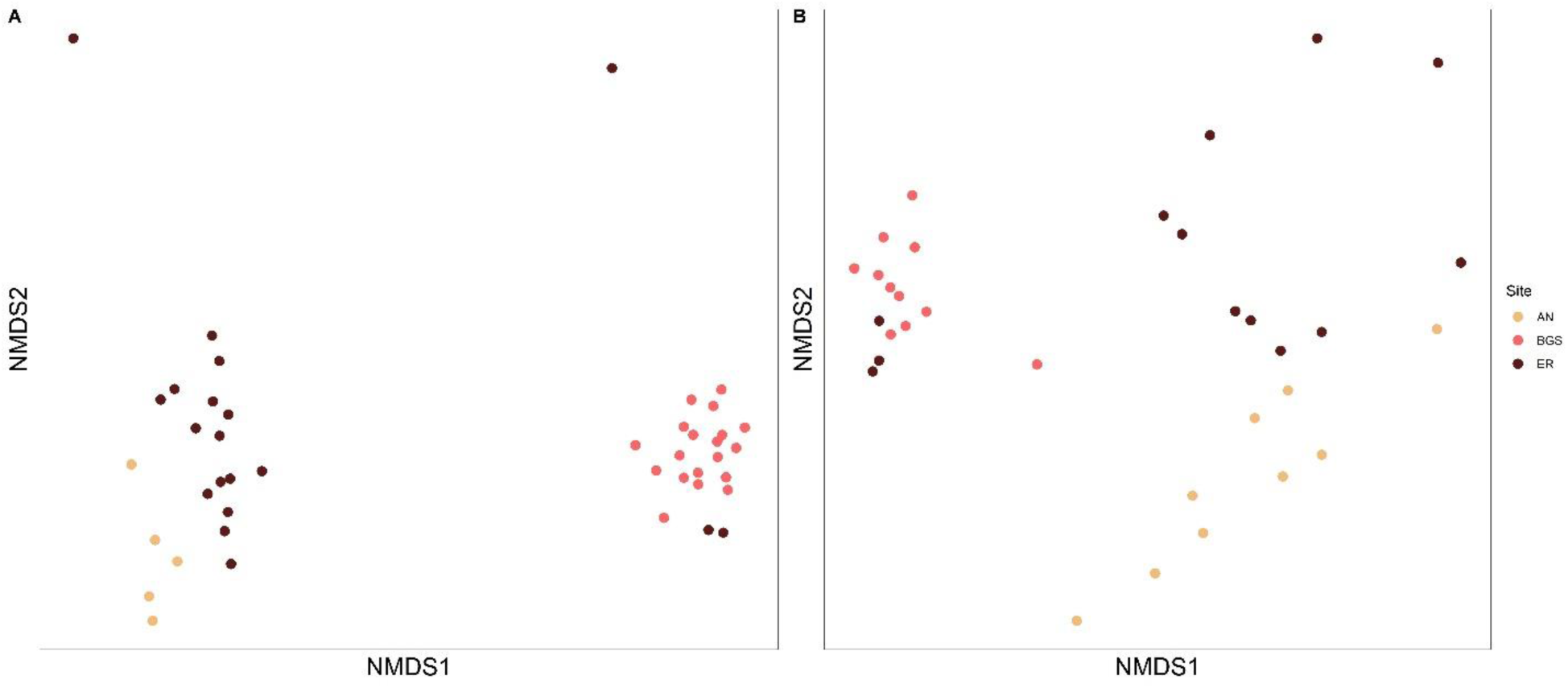
Nonmetric multidimensional scaling ordinations for the Bray-Curtis dissimilarity matrix, demonstrating clear separation in the composition of polar metabolic profile. Eurasian Blackcap (A) and Lesser Whitethroat (B) in the three different stopover sites. Freshly landed vs. rested Eurasian Blackcap (C) and Lesser Whitethroat (D).

SIMPER analysis identified ten metabolites contributing most to the dissimilarity among sites. These metabolites were identical in both warbler species and appeared significantly different at all inter-site pairwise comparisons. In Eurasian Blackcap, these metabolites contributed 48.5% (ER vs. BGS), 46.2% (AN vs. BGS), and 51.6% (AN vs. ER) to the inter-site dissimilarity. In Lesser Whitethroat, these metabolites contributed 49.6% (ER vs. BGS), 48.1% (AN vs. BGS), and 49.9% (AN vs. ER) to the inter-site dissimilarity. Among these ten metabolites, we annotated six metabolites: lactic acid, malic acid, glycerol, glycerol 3-phosphate, glucose and alanine (Fig. 2). The level of these metabolites varied significantly among sites (approx. F_12,66_=4.615, P<0.001, and approx. F_12,46_=6.575, P<0.001, for Eurasian Blackcap and Lesser Whitethroat, respectively; Tables 1S and 2S, Supplementary material). The intensity of these metabolites did not vary significantly as a function of body condition (approx. F_6,32_=0.734, P=0.626, and F_6,22_=1.897 P=0.127, for Eurasian Blackcap and Lesser Whitethroat, respectively), Nevertheless, the interaction between site and body condition was significant for Lesser Whitethroat (approx. F_12,46_=2.253 P=0.024) but not for Eurasian Blackcap (approx. F_12,66_=0.944, P=0.510). In Eurasian Blackcap, the levels of all six metabolites were significantly higher in BGS than in ER and AN (Tukey HSD p<0.01), while in Lesser Whitethroat, the level of lactic acid, alanine and glycerol 3-phosphate, were significantly higher in BGS than in ER and AN. A similar pattern was evident for malic acid with significant differences only between BGS and AN (P=0.006), glycerol with all pairwise comparisons being significant (P<0.05), and glucose with significant differences only between BGS and ER (P=0.003).

**Figure 2:**
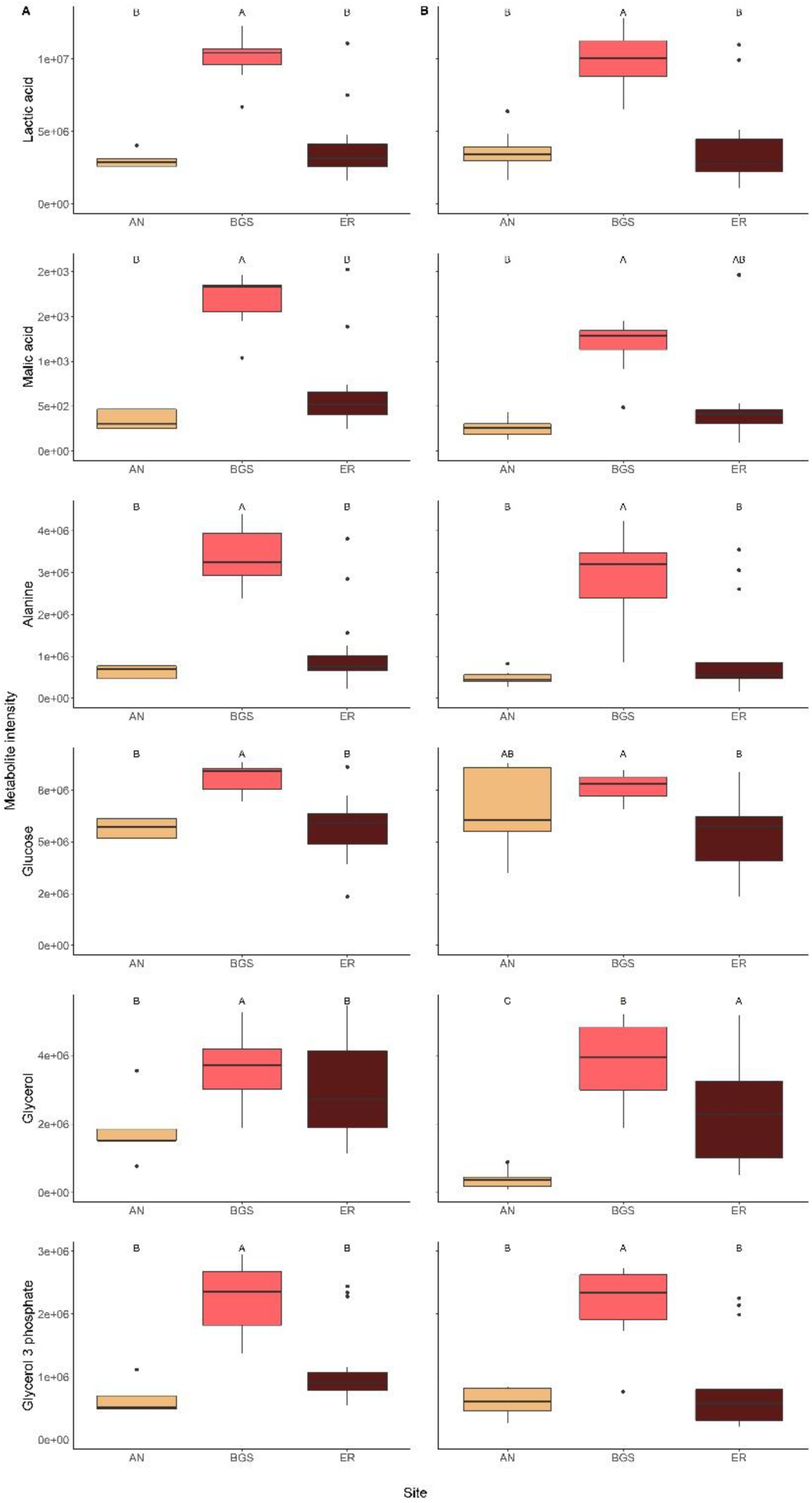
Differences in key polar metabolites in Eurasian Blackcap (A) and Lesser Whitethroat (B) among the three different stopover sites, as detected in SIMPER analyses. Different letters account for significant differences. Within boxes, horizontal lines indicate the median; black dots show the mean; box boundaries indicate the interquartile range; whiskers indicate minimum and maximum.

### Amino acids

Multivariate analysis of variance (MANOVA) followed by univariate tests, indicated that the intensities of all 12 annotated plasma amino acids were significantly higher in BGS, compared with the other two sites (approx. F_26,52_=3.291, P<0.001 and approx. F_26,30_=2.486, P=0.008, for Blackcap and Lesser Whitethroat, respectively; tables 3S and 4S, supplementary material). Additionally, individuals’ body condition did not significantly affect the plasma amino acids of both, Eurasian Blackcaps (approx. F_13,25_=2.077, P=0.057) and Lesser Whitethroats (F_13,14_=0.708, P=0.729), though the trend for Blackcaps is only marginally insignificant. Lastly, the interaction between site and body condition was not significant for both species (approx. F_26,52_=0.940, P=0.557 and approx. F_26,30_=1.379, P=0.197, for Blackcap and Lesser Whitethroat, respectively)

### Lipophilic profile

An nMDS ordination of the Bray-Curtis dissimilarity matrix, followed by a PERMANOVA indicated that in both warbler species the composition of lipophilic metabolites varied significantly among sites (Fig. 1S, supplementary material; PERMANOVAs: F_2,37_=6.6465, P>0.001, R^2^=0.237 in Eurasian Blackcap, and F_2,27_=3.540, P=0.001, R^2^=0.183 in Lesser Whitethroat). Additionally, individuals’ body condition, derived from the residuals of regressing body mass against wing length, significantly affected the lipophilic profile of both species (F_1,37_=2.523, P=0.038, R^2^=0.041 in Eurasian Blackcap, and F_1,27_=2.951, P=0.017, R^2^=0.076 in Lesser Whitethroat). The interaction between site and body condition was not significant in both species (F_2,37_=1.601, P=0.114, R^2^=0.057 in Eurasian Blackcap, and F_2,27_=0.832, P=582, R^2^=0.043 in Lesser Whitethroat). Pairwise comparisons revealed that the lipophilic profile of Blackcaps was significantly different when staging at ER compared with AN and BGS (P=0.009, and P=0.003 for Eurasian Blackcap and Lesser Whitethroat, respectively). Similarly, for Lesser whitethroats, pairwise comparison revealed significantly different lipophilic profile when staging at AN compared with ER and BGS (P=0.003 and P=0.036 for Eurasian Blackcap and Lesser Whitethroat, respectively).

SIMPER analysis identified ten metabolites contributing most to the dissimilarity among sites. In both species, and at all inter-site pairwise comparisons This list of lipids was comprised of 6-7 TAGs (50-54 carbons, with varying saturation levels of 1-5 double bonds) and 2-4 phosphatidylcholine (34-38 carbons, with varying saturation levels of 1-4 double bonds). In Eurasian Blackcap, these lipids contributed 31.8% (ER vs. BGS), 32.3% (AN vs. BGS), and 31.7% (AN vs. ER) to the inter-site dissimilarity. In Lesser Whitethroat, these metabolites contributed 33.6% (ER vs. BGS), 31.4% (AN vs. BGS), and 36.2% (AN vs. ER) to the inter-site dissimilarity. The annotated lipids mean intensities varied, and consistent pattern across sites could not be detected. We therefore added additional analyses of TAGs and BUTY.

To further examine the plasma lipids, we quantified the accumulated level of plasma TAGs (Fig. 3), manifested as intensities. Total TAG intensities were not significantly different among sites, for both species (F_2,37_=0.995, P=0.379 and F_2.26_=0.689, P=0.511, for Eurasian Blackcap and Lesser Whitethroat, respectively). Body condition did not significantly affect the total TAG intensities for both species (F_1,37_=2.050, P=0.161 and F_1,26_=0.044, P=0.836, for Eurasian Blackcap and Lesser Whitethroat, respectively). We further compared PUFA TAGs (with 6-8 double bonds within the TAG) to explore potential differences not exposed by total TAG comparison. The patterns of PUFA TAGs were consistent between the two warbler species: PUFA TAG varied among sites (F_2,37_=6.122, P=0.005 and F_2,26_=3.163, P=0.059, for Eurasian Blackcap and Lesser Whitethroat, respectively), though the trend for Lesser whitethroat could be considered as marginally insignificant. However, there was no effect of body condition on the TAGs intensity (F_1,37_=0.052, P=0.820 and F_1,26_=0.208, P=0.652, for Eurasian Blackcap and Lesser Whitethroat, respectively). Specifically, PUFA TAG intensities were higher in ER than in AN (t_37_=2.16, P=0.031, and t_26_=2.33, P=0.020 for Eurasian Blackcap and Lesser Whitethroat, respectively) and BGS, with the latter being marginally non-significant in Lesser Whitethroat (t_37_=2.16, P=0.031, and t_26_=1.7, P=0.088 for Eurasian Blackcap and Lesser Whitethroat, respectively). No significant differences in PUFA TAG intensity were detected between BGS and AN (t_37_=0.094, P=0.926, and t_26_=0.655, P=0.518 for Eurasian Blackcap and Lesser Whitethroat, respectively).

**Figure 3:**
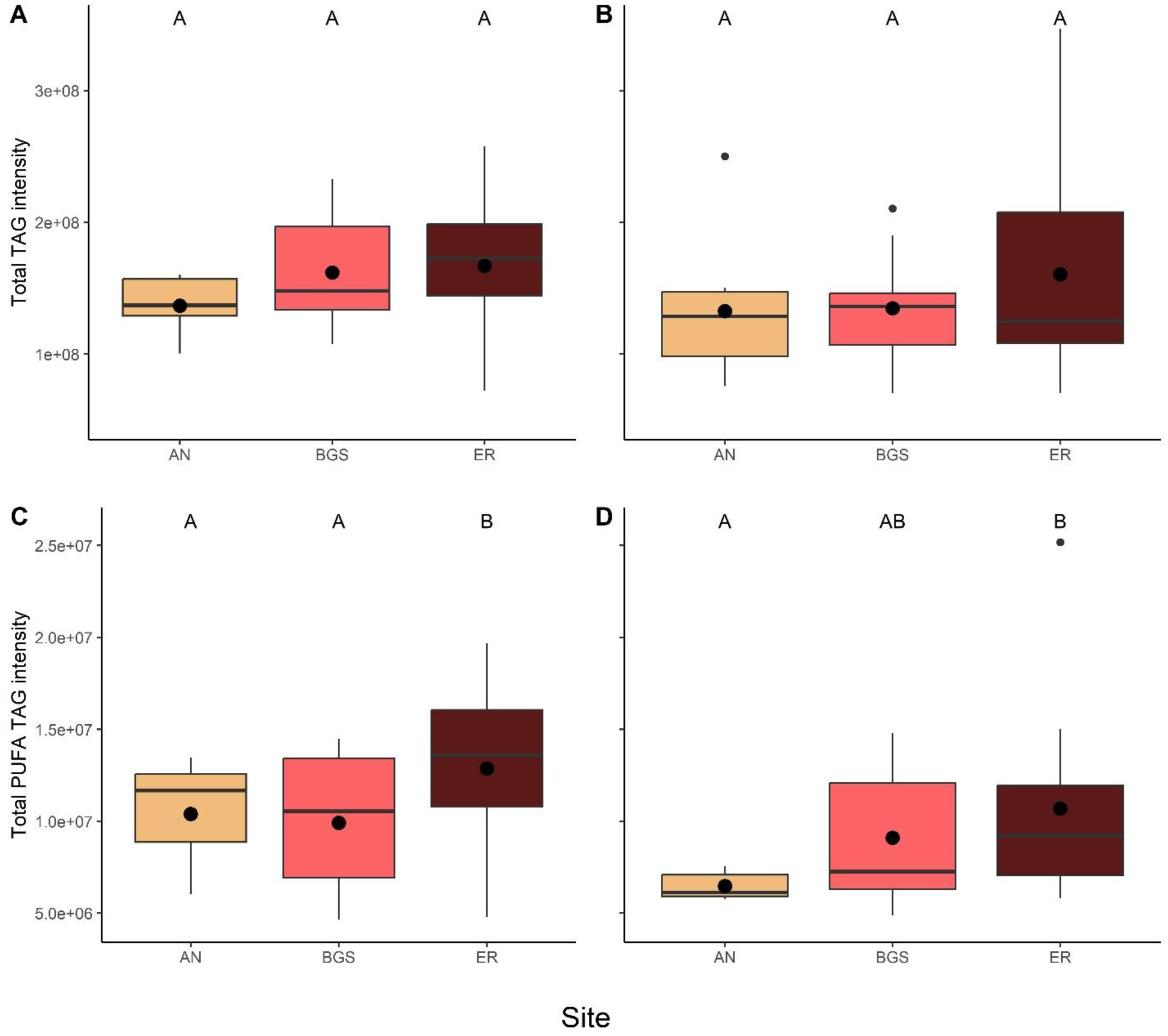
Differences in relative intensity of total TAG in Eurasian Blackcap (A) and Lesser Whitethroat (B), and total PUFA TAG in Eurasian Blackcap (C) and Lesser Whitethroat (D) among the three different stopover sites. Different letters account for significant differences. Within boxes, horizontal lines indicate the median; black dots show the mean; box boundaries indicate the interquartile range; whiskers indicate minimum and maximum.

### β-Hydroxybutyric acid

We focused on an additional metabolite, β-Hydroxybutyric acid (BUTY), which was not included in the list of metabolites detected by SIMPER but is considered to play a key role in avian metabolism (Jenni-Eiermann and Jenni 1991, Guglielmo et al. 2005), particularly during fat accumulation and ketogenesis. In both species, BUTY levels varied among sites (F_2,37_=3.480, P=0.041, and F_2,26_ = 8.304, P=0.002, for Eurasian Blackcap and Lesser Whitethroat, respectively; Fig. 4). Moreover, BUTY levels were significantly lower in AN compared with ER and BGS for both the Blackcaps (t_37_ = 2.330, P=0.025 and t_37_=2.911, P=0.006, for comparing AN with ER and BGS, respectively) and the Lesser Whitethroats (t_26_ = 2.508, P=0.019 and t_26_=3.630, P=0.001, for comparing AN with ER and BGS, respectively). Body condition significantly affected the plasma BUTY levels of the Blackcaps (F_1,37_=4.580, P=0.039) with the interaction between body condition and site also being significant (F_2,37_ = 5.044, P=0.039), however, body condition did not affect plasma BUTY levels of the lesser whitethroats (F_2,27_ = 0.067, P=0.798).

**Figure 4:**
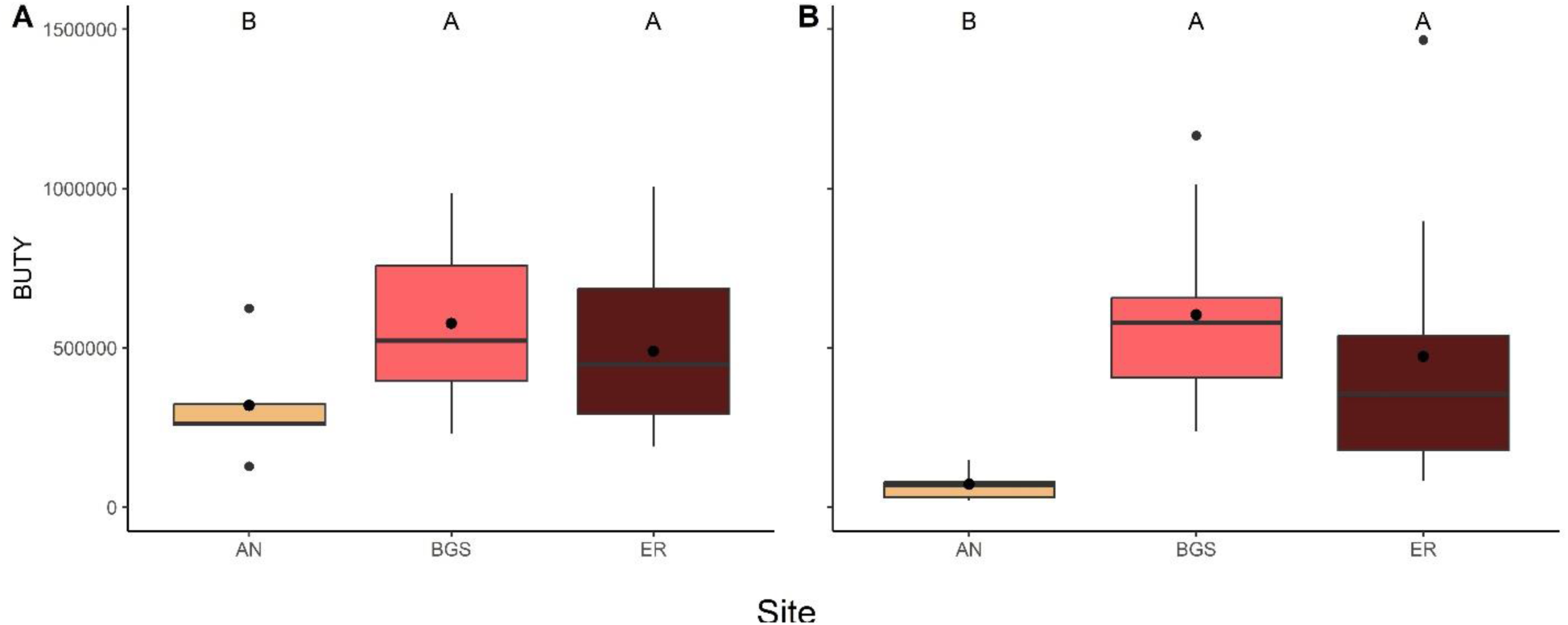
Differences in relative intensity of β-Hydroxybutyric acid (BUTY) in Eurasian Blackcap (A) and Lesser Whitethroat (B) among the three different stopover sites. Different letters account for significant differences. Within boxes, horizontal lines indicate the median; black dots show the mean; box boundaries indicate the interquartile range; whiskers indicate minimum and maximum.

## Discussion

We conducted a comparative field study to quantify the plasma metabolome of two common migratory passerine species at three different stopover sites in the northern Negev desert of Israel during autumn migration. We found that both warbler species’ polar and lipophilic metabolites varied significantly among sites. The inter-site variation in the polar metabolites can be mainly attributed to the inter-site variation in the birds’ physiological state. That is, the above-mentioned metabolites, differentiating among sites are mainly related to fasting and flight recovery. In this way, lactic acid, glucose, and glycerol are examples of metabolites that were previously demonstrated to vary between birds before and after resting (Viswanathan et al. 1987, Jenni-Eiermann and Jenni 1992). Our previous research efforts (Domer et al. 2018, 2021) have shown that during autumn migration, both recapture and fuel accumulation rates are higher in ER than in BGS. These findings, in combination with the results presented here, strongly suggest that most birds at BGS leave soon after arrival (i.e., do not spend another night at this site) while most birds in ER are at a resting/refueling state. Importantly, we could not detect significant correlation of body condition with the annotated metabolites identified by SIMPER, as well as with the amino acids, except for two distinct cases, glycerol and isoleucine, both were significantly different across body condition only for Blackcaps, with the latter also showing a significant site by body condition interaction. The inter-site variation in the lipophilic profiles of birds was harder to interpret and is suspected to reflect the variation in the primary food source.

The body condition of the birds seemed to significantly affect the plasma metabolic profile or the levels of metabolites only in distinct occasions and not for both species. That is, while body condition may have affected the plasma level of some metabolites, the variation in plasma polar metabolites is mainly associated with differences in site characteristics. Importantly, all birds captured at the arid site (BGS) were in a physiological state indicating only a short rest after flight (e.g., high lactic acid) and are suspected to have landed during the previous night. Lastly, the inter-site variation in the polar metabolic profile was mainly generated by ten metabolites, six of which were successfully annotated. Below, we discuss the involvement of these six metabolites in critical metabolic pathways activated during stopovers.

### Stopover metabolism

The polar metabolites found to vary among stopover sites were identical in both species. These metabolites mainly participate in four energy metabolism pathways: (1) fatty acid oxidation, (2) protein catabolism, (3) glucose-alanine (Cahill) cycle, and (4) lactic acid (Cori) cycle. These pathways, activated during fasting and endurance exercise, often operate simultaneously during migration. The primary energy source for long flights is derived from subcutaneous lipids. TAG degradation in the cytosol produces glycerol and glycerol 3-phosphate, which can also be a precursor for gluconeogenesis (Robergs and Griffin 1998). Fat stores are essential for birds (Pond 1978, Guglielmo 2010), as they do not carry large glycogen stores, probably due to the high cost of maintaining such hygroscopic storage molecule (Hickey et al. 2012).

In addition to glycerol, another energy source can be amino acids, derived from protein catabolism. Such catabolism occurs during flight and starvation in flight muscles, but also in the liver and other digestive organs (Bauchinger and McWilliams 2010). We found that the intensities of plasma amino acids are higher in birds that landed at BGS, which are suspected to have landed during the previous night, compared to the other sites accompanied by elevated plasma glucose. Previous research has thoroughly discussed the role protein catabolism plays during migration flight and fasting (Bauchinger and McWilliams 2012). Given that all suggested hypotheses are not mutually exclusive, and in the light of the high intensities of plasma amino acids in birds from BGS, which are likely to have landed a few hours prior capture, we suggest that an additional main pathway for these amino acids is to serve as precursors for cycling glucose in the liver via gluconeogenesis (Fig. 5).

**Figure 5:**
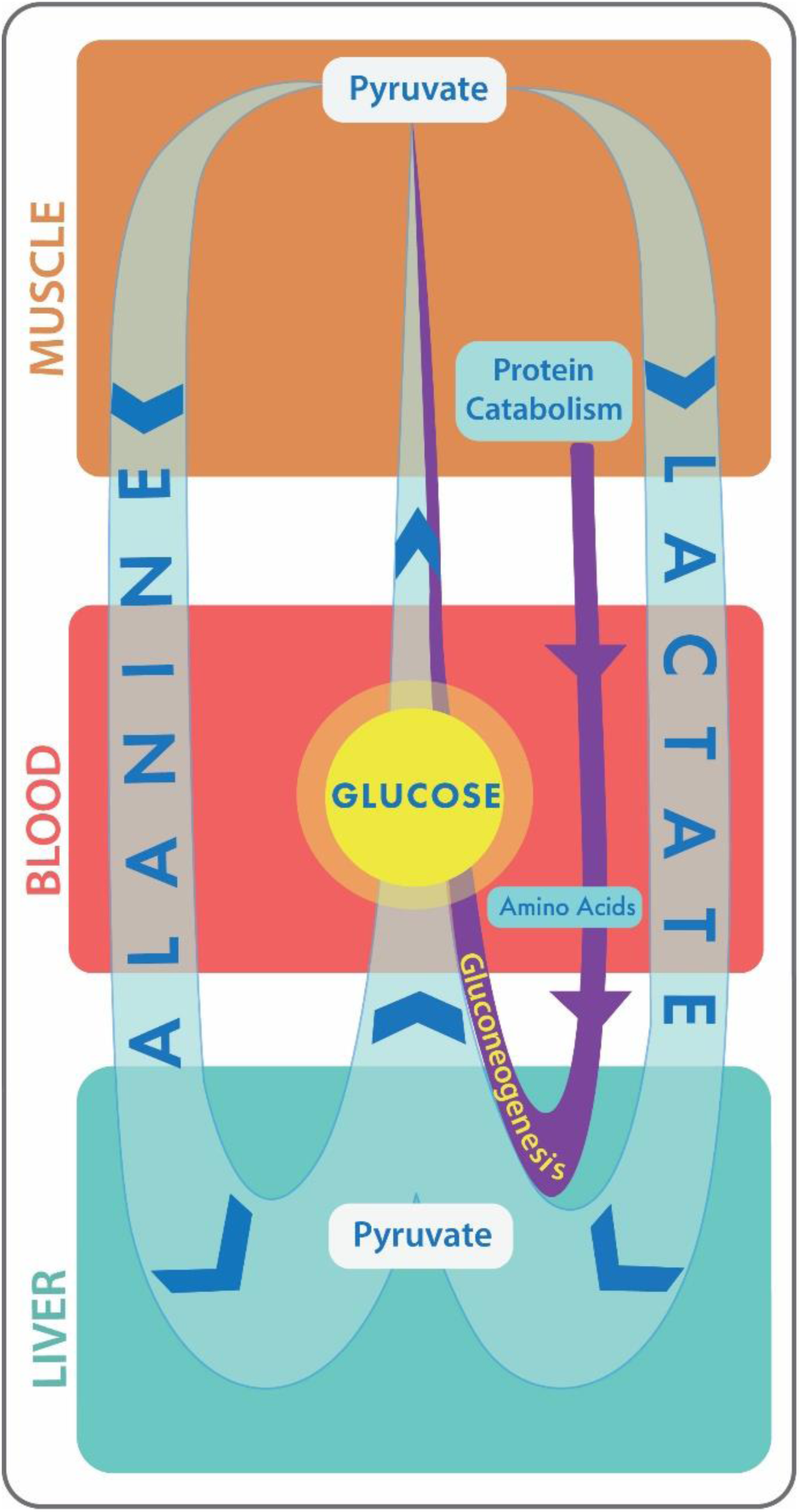
The suggested fate of protein catabolism and elevated plasma glucose during and post long-endurance flights. Free amino acids are delivered to the liver through the bloodstream. These amino acids are then used to produce glucose using gluconeogenesis. Lactic acid is maintained as a result of anaerobic conditions. The alanine cycle is maintained for disposal of the ammonium group through the uric acid cycle. The lack of NAD+ is compensated via the malate and glycerol shuttles. High plasma glucose can also facilitate repair mechanisms for high oxidative stress.

During fasting, peripheral organs become more catabolic, and such protein catabolism can support stress and healing processes by cycling glucose towards Cahill and Cori cycles (Deutz et al. 1992, Soeters and Soeters 2012). Furthermore, the cycled glucose can also facilitate reducing equivalent NADPH, which is necessary to maintain redox potential (Levin et al. 2017), a common result of endurance exercise. We, therefore, suggest that protein degradation facilitates the metabolic cycling of glucose to support physiological stress.

Cahill (alanine) and Cori (lactic acid) cycles are responsible for cycling nutrients between the skeletal muscles and liver. In the Cori cycle, the lactate, produced by anaerobic glycolysis in muscles, is transported to the liver and converted to glucose, then returns to the muscles and metabolized back to lactate, preventing the accumulation of blood lactate. The contribution of lactate to overall glucose production increases with fasting duration (Katz and Tayek 1998). Nonetheless, fasting requires utilizing substrates already present in the body. For birds, subcutaneous lipids can provide most energy for long-distance migration (Pond 1978), yet this metabolic pathway occurs alongside protein catabolism (McWilliams and Karasov 2005). In the Cahill (or alanine) cycle, the nitrogen, generated from amon acid degradation is trans-aminated to pyruvate, forming alanine (Felig 1973), and mobilized to the liver for nitrogen disposal via the urea or uric acid cycle in birds (Milroy 1903). In contrast to the Cori cycle, this pathway causes NAD^+^ deficiency, which in turn can be counteracted via the malate shuttle (Mettler and Beevers 1980) or the glycerol-3-phosphate shuttle (Shen et al. 2006). Both are mechanisms for generating NAD^+^, and are supported by our data, namely, the higher plasma malic acid, glycerol, and glycerol 3-phosphate, detected in non-rested birds.

Insulin resistance and hyperglycemia are one of the most important mechanisms for coping with prolonged fasting in animals (Soeters and Soeters 2012). Remarkably, the Cori and Cahill cycles were previously related to insulin resistance (Katz and Tayek 1998, Sarabhai and Roden 2019). Soeters et al. (2021) suggested that cycling glucose metabolites, alongside insulin resistance, are metabolically connected, serving as a beneficial survival response. They also suggested that this pattern leads to fatty acid oxidation and may be a consequence rather than a cause of insulin resistance. Adaptive insulin resistance was previously documented in some animal species, as an adaptation for living in nutrient-limited environments (Houser et al. 2013, Riddle et al. 2018). As flying vertebrates, characterized by extremely high metabolic rates, migrating birds should constantly deal with endurance exercise, even during simple movements, as well as prolonged fasting associated with migration. We suggest that avian insulin resistance and hyperglycemia are mechanisms for recovering from long-endurance flights, despite incapability of feeding.

### Ecological perspective

Here we show that the same ten polar metabolites, which largely generate the inter-site dissimilarly in the metabolome of both warbler species, are highly related to the physiological status of the birds, suggesting they have recently landed from long flight. While the detected inter-site variation in the polar metabolic profile could be attributed to variation in the birds’ physiological state, this intrinsic factor could not explain the observed inter-site variation in the lipophilic profiles. Most birds in ER and AN were in a resting/refueling state, although these sites offered them different food types (fat-rich fruits and nectar, respectively). Nevertheless, there were significant differences not only in their lipophilic profile but also in their PUFA TAGs, and BUTY intensities, which were higher in ER and are known to increase not only when fasting, but also when feeding on a lipid food source (Smith et al. 2007). We therefore suggest that the lipophilic profile variation between ER and AN should be attributed to the primary food source these two sites provide. Nectar is comprised mainly of sugars dissolved in water which are absorbed quickly into the digestive tract of birds (Tracy et al. 2010). Therefore, the blood glucose associated with nectar consumption may have little or no immediate effect on the respective lipophilic profile compared to the consumption of fat-rich fruits. (Jenni-Eiermann and Jenni 1991)

### Conclusions

Although lipid metabolism is considered as the primary metabolic pathway during long-endurance flights (Ramenofsky 1990, Jenni-Eiermann 2017), the results of blood lipid profiles were hard to interpret, as they contained many lipophilic compounds that do not necessarily relate to lipid metabolism during exercise. Additionally, TAG and BUTY levels were not a good indicator of site quality. These findings are consistent with previous research suggesting that the context of these metabolites may be species-specific or related to food sources (Jenni-Eiermann and Jenni 1992, Guglielmo et al. 2005, Smith et al. 2007). Essentially, the pathways proposed here to be activated during a stopover indicate a need for flight recovery and suggest that glucose cycling, derived from protein catabolism, plays a vital role in this recovery process. Moreover, this viewpoint also suggests that avian insulin resistance and hyperglycemia have evolved due to endurance exercise, prolonged fasting, and fatty acid oxidation, similar to trauma recovery in other animals.

## Contact information

Adi Domer –

Weronika Jensinka –

Leah Rosental –

Eyal Shochat –

Alisdair R. Fernie –

Saleh Alseekh –

Yariv Brotman –

Ofer Ovadia –

All authors declare to have no competing interests.

The data that support the findings of this study will be made openly available in a repository.

## References

Abdel-Rachied, H. G., S. A. Zaahkouk, E. I. El-Zawhry, and K. S. Elfeky. 2014. Flying or running stress effect on some hematological and biochemical parameters in some birds and mammals. J. Entomol. Zool. Stud 2:153–158.

Aidley, D. J. 1981. Animal migration. CUP Archive.

Bauchinger, U., and S. R. McWilliams. 2010. Extent of phenotypic flexibility during long-distance flight is determined by tissue-specific turnover rates: a new hypothesis. Journal of Avian Biology 41:603–608.

Bauchinger, U., and S. R. McWilliams. 2012. Tissue-specific mass changes during fasting: the protein turnover hypothesis. Pages 193–206 Comparative physiology of fasting, starvation, and food limitation. Springer.

Berthold, P. 1996. Control of bird migration. Springer Science & Business Media.

Blem, C. R. 1976. Patterns of lipid storage and utilization in birds. Integrative and Comparative Biology 16:671–684.

Bordel, R., and E. Haase. 1993. Effects of flight on blood parameters in homing pigeons. Journal of Comparative Physiology B 163:219–224.

Braun, E. J., and K. L. Sweazea. 2008. Glucose regulation in birds. Comparative Biochemistry and Physiology Part B: Biochemistry and Molecular Biology 151:1–9.

Clarke, A., and H. Pörtner. 2010. Temperature, metabolic power and the evolution of endothermy. Biological Reviews 85:703–727.

Clarke, A., and P. Rothery. 2008. Scaling of body temperature in mammals and birds. Functional Ecology 22:58–67.

Costantini, D., M. Cardinale, and C. Carere. 2007. Oxidative damage and anti-oxidant capacity in two migratory bird species at a stop-over site. Comparative Biochemistry and Physiology Part C: Toxicology & Pharmacology 144:363–371.

Cuadros-Inostroza, A., C. Caldana, H. Redestig, M. Kusano, J. Lisec, H. Peña-Cortés, L. Willmitzer, and M. A. Hannah. 2009. TargetSearch-a Bioconductor package for the efficient preprocessing of GC-MS metabolite profiling data. BMC bioinformatics 10:1–12.

Dai, C., W. Zhang, R. He, F. Xiong, and H. Ma. 2017. Protein breakdown and release of antioxidant peptides during simulated gastrointestinal digestion and the absorption by everted intestinal sac of rapeseed proteins. LWT 86:424–429.

Deutz, N. E. P., P. L. M. Reijven, G. Athanasas, and P. B. Soeters. 1992. Post-operative changes in hepatic, intestinal, splenic and muscle fluxes of amino acids and ammonia in pigs. Clinical Science 83:607–614.

Domer, A., O. Ovadia, and E. Shochat. 2018. Energy for the road: Influence of carbohydrate and water availability on fueling processes in autumn-migrating passerines. Auk 135.

Domer, A., E. Vinepinsky, A. Bouskila, E. Shochat, and O. Ovadia. 2021. Optimal Stopover Model: a state dependent habitat selection model for staging passerines. Journal of Animal Ecology.

Domer Adi. 2023. Comparative analysis of the plasma metabolome of migrating passerines during stopover: Novel insights into flight metabolism, Dryad, Dataset, 10.5061/dryad.k3j9kd5cf.

Faraci, F. M. 1991. Adaptations to hypoxia in birds: how to fly high. Annual Review of Physiology 53:59–70.

Felig, P. 1973. The glucose-alanine cycle. Metabolism 22:179–207.

Gerson, A. R., and C. G. Guglielmo. 2011. Flight at low ambient humidity increases protein catabolism in migratory birds. Science 333:1434–1436.

Gómez, C., N. J. Bayly, D. R. Norris, S. A. Mackenzie, K. V Rosenberg, P. D. Taylor, K. A. Hobson, and C. D. Cadena. 2017. Fuel loads acquired at a stopover site influence the pace of intercontinental migration in a boreal songbird. Scientific reports 7:1–11.

Guglielmo, C. G. 2010. Move that fatty acid: Fuel selection and transport in migratory birds and bats. Integrative and Comparative Biology 50:336–345.

Guglielmo, C. G., D. J. Cerasale, and C. Eldermire. 2005. A field validation of plasma metabolite profiling to assess refueling performance of migratory birds. Physiological and Biochemical Zoology 78:116–125.

Hickey, A. J. R., M. Jüllig, J. Aitken, K. Loomes, M. E. Hauber, and A. R. J. Phillips. 2012. Birds and longevity: does flight driven aerobicity provide an oxidative sink? Ageing research reviews 11:242–253.

Houser, D. S., C. D. Champagne, and D. E. Crocker. 2013. A non-traditional model of the metabolic syndrome: the adaptive significance of insulin resistance in fasting-adapted seals. Frontiers in endocrinology 4:164.

Hummel, J., S. Segu, Y. Li, S. Irgang, J. Jueppner, and P. Giavalisco. 2011. Ultra performance liquid chromatography and high resolution mass spectrometry for the analysis of plant lipids. Frontiers in plant science 2:54.

Jenni, L., and S. Jenni-Eiermann. 1998. Fuel supply and metabolic constraints in migrating birds. Journal of Avian Biology:521–528.

Jenni-Eiermann, S. 2017. Energy metabolism during endurance flight and the post-flight recovery phase. Journal of Comparative Physiology A 203:431–438.

Jenni-Eiermann, S., and L. Jenni. 1991. Metabolic responses to flight and fasting in night-migrating passerines. Journal of Comparative Physiology B 161:465–474.

Jenni-Eiermann, S., and L. Jenni. 1992. High plasma triglyceride levels in small birds during migratory flight: a new pathway for fuel supply during endurance locomotion at very high mass-specific metabolic rates? Physiological Zoology 65:112–123.

Jenni-Eiermann, S., and L. Jenni. 2001. Postexercise ketosis in night-migrating passerine birds. Physiological and Biochemical Zoology 74:90–101.

Jenni-Eiermann, S., L. Jenni, A. Kvist, A. Lindström, T. Piersma, and G. H. Visser. 2002. Fuel use and metabolic response to endurance exercise: a wind tunnel study of a long-distance migrant shorebird. Journal of Experimental Biology 205:2453–2460.

Katz, J., and J. A. Tayek. 1998. Gluconeogenesis and the Cori cycle in 12-, 20-, and 40-h-fasted humans. American Journal of Physiology-Endocrinology And Metabolism 275:E537–E542.

Lapidot-Cohen, T., L. Rosental, and Y. Brotman. 2020. Liquid Chromatography–Mass Spectrometry (LC-MS)-Based Analysis for Lipophilic Compound Profiling in Plants. Current protocols in plant biology 5:e20109.

Levin, E., G. Lopez-Martinez, B. Fane, and G. Davidowitz. 2017. Hawkmoths use nectar sugar to reduce oxidative damage from flight. Science 355:733–735.

Luo, P., Z. Wang, C. Su, H. Li, H. Zhang, Y. Huang, and W. Chen. 2023. Chicken GLUT4 undergoes complex alternative splicing events and its expression in striated muscle changes dramatically during development. Poultry Science 102:102403.

Maillet, D., and J.-M. Weber. 2006. Performance-enhancing role of dietary fatty acids in a long-distance migrant shorebird: the semipalmated sandpiper. Journal of Experimental Biology 209:2686–2695.

McWilliams, S. R., and W. H. Karasov. 2001. Phenotypic flexibility in digestive system structure and function in migratory birds and its ecological significance. Comparative Biochemistry and Physiology Part A: Molecular & Integrative Physiology 128:577–591.

McWilliams, S. R., and W. H. Karasov. 2005. Migration takes guts. Birds of two worlds: the ecology and evolution of migration. Smithsonian Institution Press, Washington, DC:67–78.

Mettler, I. J., and H. Beevers. 1980. Oxidation of NADH in glyoxysomes by a malate-aspartate shuttle. Plant physiology 66:555–560.

Milroy, T. H. 1903. The formation of uric acid in birds. The Journal of physiology 30:47.

Pond, C. M. 1978. Morphological aspects and the ecological and mechanical consequences of fat deposition in wild vertebrates. Annual Review of Ecology and Systematics 9:519–570.

Potter, J. H. T., R. Drinkwater, K. T. J. Davies, N. Nesi, M. C. W. Lim, L. R. Yohe, H. Chi, X. Zhang, I. Levantis, and B. K. Lim. 2021. Nectar-feeding bats and birds show parallel molecular adaptations in sugar metabolism enzymes. Current Biology 31:4667–4674.

Ramenofsky, M. 1990. Fat storage and fat metabolism in relation to migration. Pages 214–231 Bird migration. Springer.

Riddle, M. R., A. C. Aspiras, K. Gaudenz, R. Peuß, J. Y. Sung, B. Martineau, M. Peavey, A. C. Box, J. A. Tabin, and S. McGaugh. 2018. Insulin resistance in cavefish as an adaptation to a nutrient-limited environment. Nature 555:647–651.

Robergs, R. A., and S. E. Griffin. 1998. Glycerol. Sports Medicine 26:145–167.

Robin, J.-P., Y. Cherel, H. Girard, A. Géloen, and Y. Le Maho. 1987. Uric acid and urea in relation to protein catabolism in long-term fasting geese. Journal of Comparative Physiology B 157:491–499.

Sapir, N., I. Tsurim, B. Gal, and Z. Abramsky. 2004. The effect of water availability on fuel deposition of two staging Sylvia warblers. Journal of Avian Biology 35:25–32.

Sarabhai, T., and M. Roden. 2019. Hungry for your alanine: when liver depends on muscle proteolysis. The Journal of clinical investigation 129:4563–4566.

Satoh, T. 2021. Bird evolution by insulin resistance. Trends in Endocrinology & Metabolism 32:803–813.

Schauer, N., Y. Semel, I. Balbo, M. Steinfath, D. Repsilber, J. Selbig, T. Pleban, D. Zamir, and A. R. Fernie. 2008. Mode of inheritance of primary metabolic traits in tomato. The Plant Cell 20:509–523.

Schwilch, R., L. Jenni, and S. Jenni-Eiermann. 1996a. Metabolic responses of homing pigeons to flight and subsequent recovery. Journal of Comparative Physiology B 166:77–87.

Schwilch, R., L. Jenni, and S. Jenni-Eiermann. 1996b. Metabolic responses of homing pigeons to flight and subsequent recovery. Journal of Comparative Physiology B 166:77–87.

Seaman, D. A., C. G. Guglielmo, and T. D. Williams. 2005. Effects of physiological state, mass change and diet on plasma metabolite profiles in the western sandpiper Calidris mauri. Journal of Experimental Biology 208:761–769.

Shen, W., Y. Wei, M. Dauk, Y. Tan, D. C. Taylor, G. Selvaraj, and J. Zou. 2006. Involvement of a glycerol-3-phosphate dehydrogenase in modulating the NADH/NAD+ ratio provides evidence of a mitochondrial glycerol-3-phosphate shuttle in Arabidopsis. The Plant Cell 18:422–441.

Smith, S. B., S. R. McWilliams, and C. G. Guglielmo. 2007. Effect of diet composition on plasma metabolite profiles in a migratory songbird. The Condor 109:48–58.

Soeters, M. R., and P. B. Soeters. 2012. The evolutionary benefit of insulin resistance. Clinical nutrition 31:1002–1007.

Soeters, P. B., A. Shenkin, L. Sobotka, M. R. Soeters, P. W. de Leeuw, and R. R. Wolfe. 2021. The anabolic role of the Warburg, Cori-cycle and Crabtree effects in health and disease. Clinical Nutrition 40:2988–2998.

Stevens, L. 2004. Avian biochemistry and molecular biology. Cambridge university press.

Team, R. C. 2013. R: A language and environment for statistical computing.

Tracy, C. R., T. J. McWhorter, M. S. Wojciechowski, B. Pinshow, and W. H. Karasov. 2010. Carbohydrate absorption by blackcap warblers (Sylvia atricapilla) changes during migratory refuelling stopovers. Journal of Experimental Biology 213:380–385.

Viswanathan, M., T. M. John, J. C. George, and R. J. Etches. 1987. Flight effects on plasma glucose, lactate, catecholamines and corticosterone in homing pigeons. Hormone and metabolic research 19:400–402.

Wilcove, D. S., and M. Wikelski. 2008. Going, going, gone: is animal migration disappearing. PLoS biology 6:e188.

