## Supplementary material for "Comparative analysis of the plasma metabolome of migrating passerines during stopover: Novel insights into flight metabolism"

Table 1S. Mean intensity levels of annotated metabolites, detected in SIMPER analysis, in Eurasian Blackcaps and Lesser Whitethroat sampled sampled at our three stopover sites.

| | **Eurasian Blackcap** | | | **Lesser Whitethroats** | | |
|---|---|---|---|---|---|---|
| **Metabolite** | **Mean AN** | **Mean BGS** | **Mean ER** | **Mean AN** | **Mean BGS** | **Mean ER** |
| Lactic acid | 2955671 | 11376185 | 4280006 | 3533995 | 10320452 | 4759944 |
| Malic acid | 355.14 | 2024.205 | 627.8833 | 265.1902 | 1182.332 | 801.4244 |
| Alanine | 664939 | 4057549 | 1087954 | 494465.7 | 2923916 | 1089678 |
| Glycerol | 1846721 | 4624917 | 3152482 | 413651.6 | 5228089 | 2918094 |
| Glycerol-3-phosphate | 638840.1 | 2598702 | 1097823 | 612919.4 | 2551518 | 880673.3 |
| Glucose | 5632176 | 8458208 | 5887058 | 7121984 | 9025502 | 6001319 |

Table 2S. Results of univariate analysis of variance, with plasma metabolite intensity as the response variables, site as a categorical variable, and body condition (derived from the residuals of regressing individuals' body mass against wing length) as a covariate, for Eurasian Blackcaps and Lesser Whitethroat.

| | | **Eurasian Blackcap** | | **Lesser Whitethroats** | |
|---|---|---|---|---|---|
| Metabolite | Response variable | F value | P value | F value | P value |
| Lactic acid | Site | $F_{2,37}$=37.560 | **>0.001** | $F_{2,27}$=15.182 | **>0.001** |
| | Body Condition | $F_{1,37}$=0.050 | 0.824 | $F_{1,27}$=0.310 | 0.582 |
| | Site × Body Condition | $F_{2,37}$=0.036 | 0.964 | $F_{2,27}$=0.831 | 0.446 |
| Malic acid | Site | $F_{2,37}$=59.643 | **>0.001** | $F_{2,27}$=5.332 | **0.011** |
| | Body Condition | $F_{1,37}$=0.057 | 0.813 | $F_{1,27}$=0.470 | 0.49886 |
| | Site × Body Condition | $F_{2,37}$=0.474 | 0.626 | $F_{2,27}$=0.525 | 0.59737 |
| Alanine | Site | $F_{2,37}$=46.783 | **>0.001** | $F_{2,27}$=17.847 | **>0.001** |
| | Body Condition | $F_{1,37}$=0.091 | 0.765 | $F_{1,27}$=0.029 | 0.865 |
| | Site × Body Condition | $F_{2,37}$=0.060 | 0.942 | $F_{2,27}$=0.782 | 0.467 |
| Glycerol | Site | $F_{2,37}$=9.583 | **>0.001** | $F_{2,27}$=12.9129 | **>0.001** |
| | Body Condition | $F_{1,37}$=4.115 | **0.0497** | $F_{1,27}$=0.1323 | 0.719 |
| | Site × Body Condition | $F_{2,37}$=4.859 | **0.013** | $F_{2,27}$=2.2724 | 0.122 |
| Glycerol 3P | Site | $F_{2,37}$=35.409 | **>0.001** | $F_{2,27}$=24.653 | **>0.001** |
| | Body Condition | $F_{1,37}$=0.009 | 0.926 | $F_{1,27}$=2.590 | 0.119 |
| | Site × Body Condition | $F_{2,37}$=0.252 | 0.778 | $F_{2,27}$=0.912 | 0.414 |
| Glucose | Site | $F_{2,37}$=21.229 | **>0.001** | $F_{2,27}$=6.545 | **0.005** |
| | Body Condition | $F_{1,37}$=0.0484 | 0.827 | $F_{1,27}$=0.619 | 0.4381 |
| | Site × Body Condition | $F_{2,37}$=0.211 | 0.811 | $F_{2,27}$=1.589 | 0.223 |

Table 3S. Mean intensity levels of plasma amino acids in Eurasian Blackcaps and Lesser Whitethroat sampled sampled at our three stopover sites.

| | **Eurasian Blackcap** | | | **Lesser Whitethroats** | | |
|---|---|---|---|---|---|---|
| **Amino acid** | **Mean AN** | **Mean BGS** | **Mean ER** | **Mean AN** | **Mean BGS** | **Mean ER** |
| Glycine | 276377 | 1375346 | 609837.2 | 54037.78 | 376410.1 | 149107.8 |
| Proline | 199703.4 | 955606.8 | 454933.8 | 99231.22 | 514625.8 | 391491.4 |
| Serine | 156963.6 | 774254.7 | 231354.1 | 144350.9 | 724587 | 258461.9 |
| Threonine | 48829.8 | 136897.3 | 62162.47 | 19008.89 | 101226.1 | 43843.92 |
| Valine | 120854.4 | 550105.9 | 305590.5 | 141724.1 | 583777.5 | 331545.5 |
| Alanine | 769109.4 | 4623796 | 1230191 | 572731.3 | 3349246 | 1266236 |
| Aspartic acid | 7469.8 | 33397.84 | 14593.11 | 10642.22 | 54534.73 | 24905.92 |
| Glutamic acid | 44299.4 | 152265.9 | 78325.26 | 25608.44 | 130070.2 | 64830.69 |
| Isolucine | 33125.4 | 168448.9 | 75949.16 | 31102.33 | 164969.6 | 94946.38 |
| Leucin | 60544.4 | 394986.9 | 156731.2 | 61799.75 | 337619.8 | 192960.3 |
| Methionine | 15564.6 | 69863.84 | 17197.68 | 15458.56 | 70948.36 | 27286.08 |
| Ornithine | 6578.2 | 43047.11 | 15416.26 | 4192.111 | 32363.91 | 16098.23 |
| Tyrosine | 33328.6 | 226454.8 | 40996.79 | 18273.33 | 102191.7 | 44373.31 |

Table 4S. Results of univariate analysis of variance, with plasma amino acid intensity as response variables, the site as a categorical variable, and body condition (derived from the residuals of regressing individuals' body mass against wing length) as a covariate, for Eurasian Blackcaps and Lesser Whitethroat.

| | | **Eurasian Blackcap** | | **Lesser Whitethroats** | |
|---|---|---|---|---|---|
| Amino Acid | Response variable | F value | P value | F value | P value |
| Glycine | Site | $F_{2,37}$=30.064 | **>0.001** | $F_{2,26}$=16.700 | **>0.001** |
| | Body Condition | $F_{1,37}$=3.392 | 0.074 | $F_{1,26}$=0.208 | 0.652 |
| | Site × Body Condition | $F_{2,37}$=0.639 | 0.534 | $F_{2,26}$=0.476 | 0.627 |
| Proline | Site | $F_{2,37}$=17.059 | **>0.001** | $F_{2,26}$=3.067 | 0.064 |
| | Body Condition | $F_{1,37}$=0.831 | 0.368 | $F_{1,26}$=1.087 | 0.307 |
| | Site × Body Condition | $F_{2,37}$=0.563 | 0.575 | $F_{2,26}$=0.085 | 0.919 |
| Serine | Site | $F_{2,37}$=27.433 | **>0.001** | $F_{2,26}$=17.414 | **>0.001** |
| | Body Condition | $F_{1,37}$=3.227 | 0.081 | $F_{1,26}$=0.048 | 0.828 |
| | Site × Body Condition | $F_{2,37}$=0.094 | 0.910 | $F_{2,26}$=0.247 | 0.783 |
| Threonine | Site | $F_{2,37}$=9.831 | **>0.001** | $F_{2,26}$=15.630 | **>0.001** |
| | Body Condition | $F_{1,37}$=1.072 | 0.307 | $F_{1,26}$=0.175 | 0.679 |
| | Site × Body Condition | $F_{2,37}$=0.274 | 0.762 | $F_{2,26}$=0.248 | 0.782 |
| Valine | Site | $F_{2,37}$=19.726 | 0.000 | $F_{2,26}$=8.531 | **0.001** |
| | Body Condition | $F_{1,37}$=1.835 | 0.184 | $F_{1,26}$=0.213 | 0.649 |
| | Site × Body Condition | $F_{2,37}$=1.140 | 0.331 | $F_{2,26}$=0.418 | 0.663 |
| Alanine | Site | $F_{2,37}$=47.548 | **>0.001** | $F_{2,26}$=18.647 | **>0.001** |
| | Body Condition | $F_{1,37}$=0.116 | 0.735 | $F_{1,26}$=0.295 | 0.592 |
| | Site × Body Condition | $F_{2,37}$=0.094 | 0.911 | $F_{2,26}$=0.284 | 0.755 |
| Aspartic acid | Site | $F_{2,37}$=10.896 | **>0.001** | $F_{2,26}$=9.808 | **0.001** |
| | Body Condition | $F_{1,37}$=0.125 | 0.725 | $F_{1,26}$=0.683 | 0.416 |
| | Site × Body Condition | $F_{2,37}$=0.496 | 0.613 | $F_{2,26}$=0.023 | 0.977 |
| Glutamic acid | Site | $F_{2,37}$=4.745 | 0.015 | $F_{2,26}$=11.449 | **>0.001** |
| | Body Condition | $F_{1,37}$=2.425 | 0.128 | $F_{1,26}$=0.898 | 0.352 |
| | Site × Body Condition | $F_{2,37}$=0.439 | 0.648 | $F_{2,26}$=0.156 | 0.856 |
| Isolucine | Site | $F_{2,37}$=22.198 | **>0.001** | $F_{2,26}$=9.234 | **0.001** |
| | Body Condition | $F_{1,37}$=10.476 | **0.003** | $F_{1,26}$=0.162 | 0.691 |
| | Site × Body Condition | $F_{2,37}$=4.911 | **0.013** | $F_{2,26}$=1.331 | 0.282 |
| Leucin | Site | $F_{2,37}$=38.565 | **>0.001** | $F_{2,26}$=11.774 | **>0.001** |
| | Body Condition | $F_{1,37}$=2.891 | 0.097 | $F_{1,26}$=0.356 | 0.556 |
| | Site × Body Condition | $F_{2,37}$=4.862 | 0.013 | $F_{2,26}$=1.358 | 0.275 |
| Methionine | Site | $F_{2,37}$=44.914 | **>0.001** | $F_{2,26}$=11.716 | **>0.001** |
| | Body Condition | $F_{1,37}$=2.321 | 0.136 | $F_{1,26}$=0.995 | 0.328 |
| | Site × Body Condition | $F_{2,37}$=0.321 | 0.728 | $F_{2,26}$=0.064 | 0.938 |
| Ornithine | Site | $F_{2,37}$=32.965 | **>0.001** | $F_{2,26}$=9.510 | **0.001** |
| | Body Condition | $F_{1,37}$=1.814 | 0.186 | $F_{1,26}$=0.927 | 0.344 |
| | Site × Body Condition | $F_{2,37}$=1.185 | 0.317 | $F_{2,26}$=0.189 | 0.829 |
| Tyrosine | Site | $F_{2,37}$=45.336 | **>0.001** | $F_{2,26}$=12.252 | **>0.001** |
| | Body Condition | $F_{1,37}$=0.510 | 0.480 | $F_{1,26}$=0.547 | 0.466 |
| | Site × Body Condition | $F_{2,37}$=0.639 | 0.534 | $F_{2,26}$=0.314 | 0.733 |

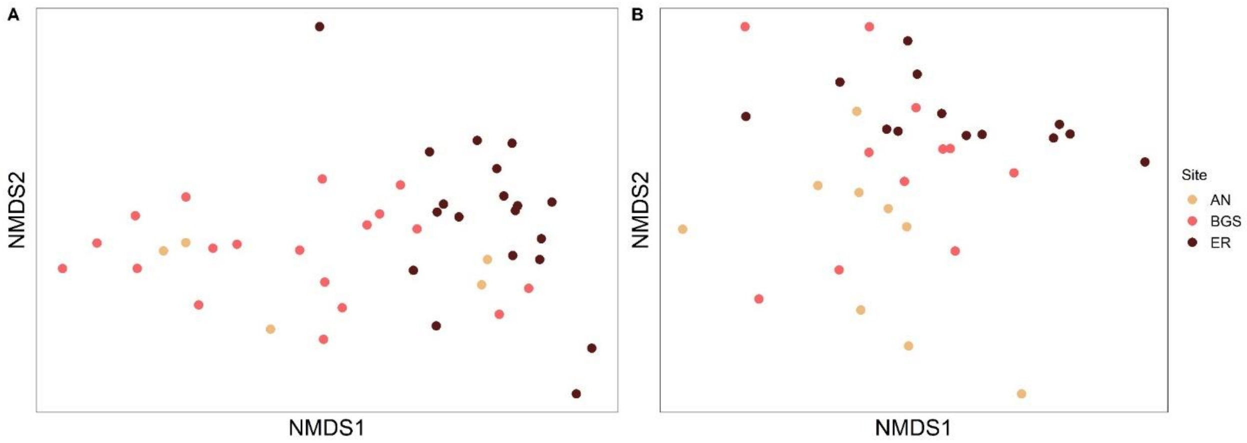


Figure 1S: Figure 1: Nonmetric multidimensional scaling ordinations for the Bray-Curtis dissimilarity matrix, exploring for differences in the lipophilic profiles. Eurasian Blackcap (A) and Lesser Whitethroat (B) in the three different stopover sites.

Optional for supplementary

Dispersion BC lipids

LW lipids

Dispersion

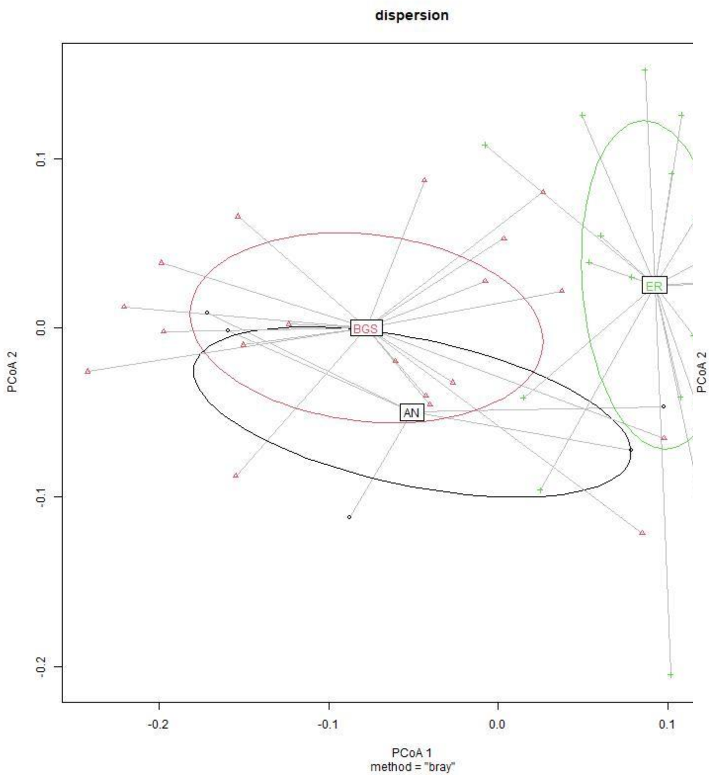


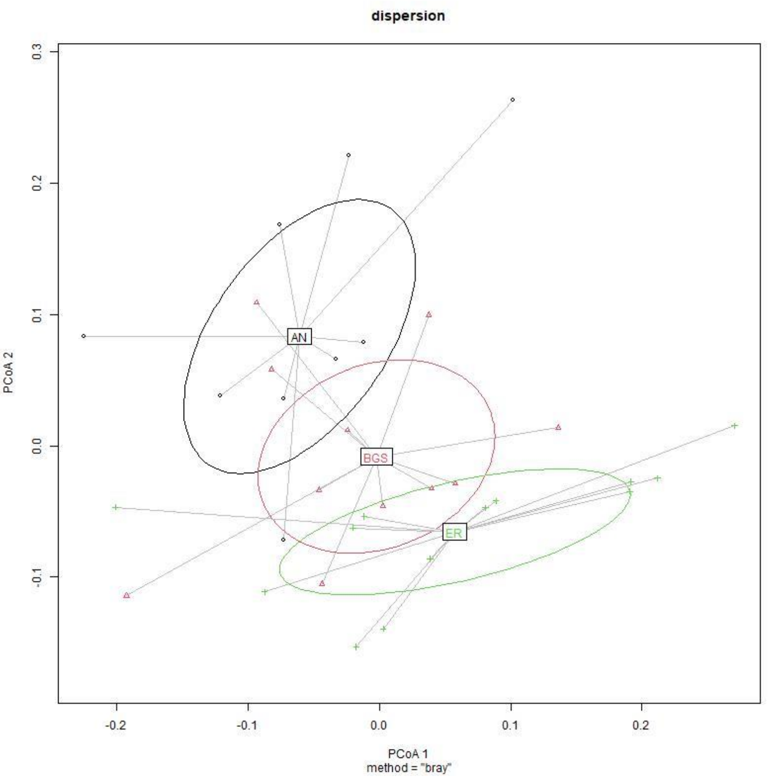

Dispersion BC polar

LW polar

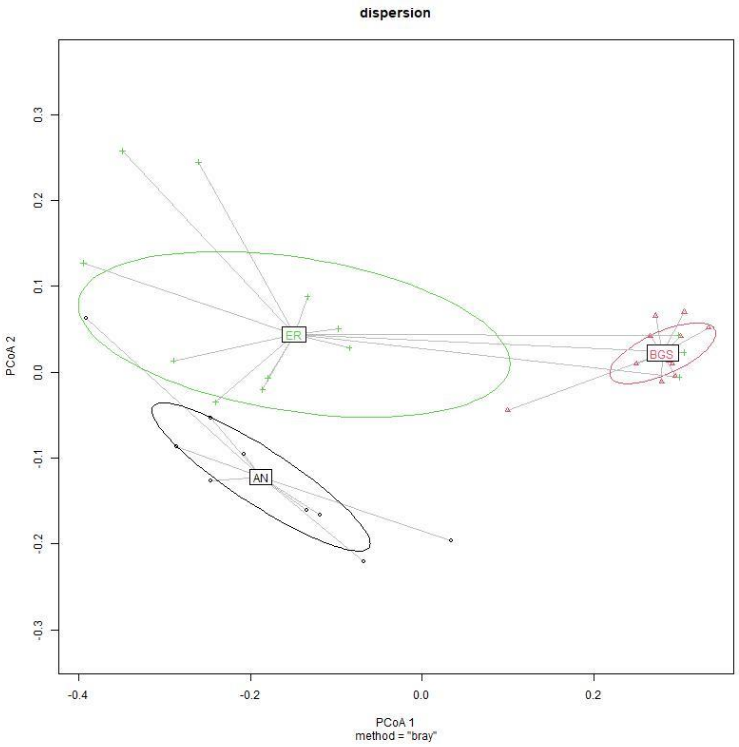


Dispersion

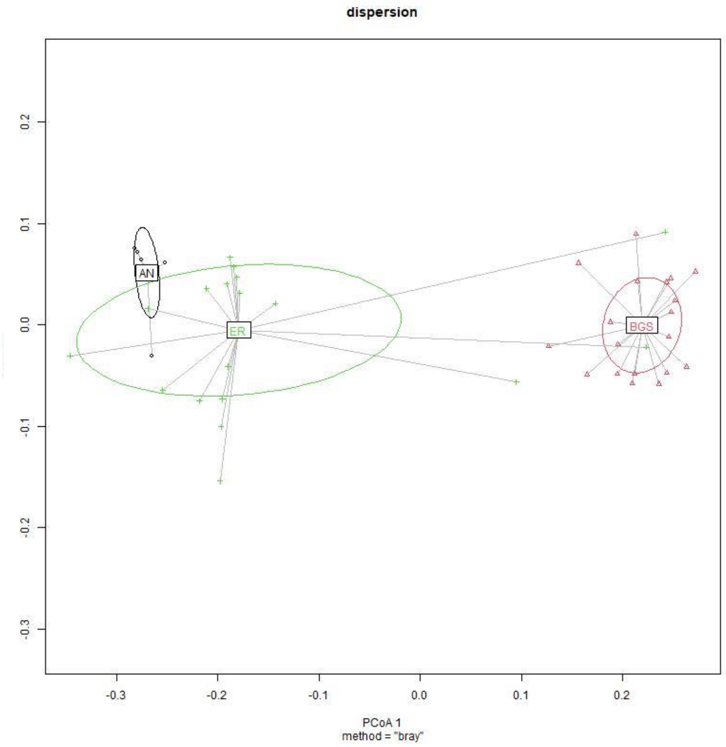
